# A meta-analysis of global avian survival across species and latitude

**DOI:** 10.1101/805705

**Authors:** Micah N. Scholer, Matt Strimas-Mackey, Jill E. Jankowski

## Abstract

Tropical birds are purported to be longer lived than temperate species of similar size, but it has not been shown whether avian survival rates covary with a latitudinal gradient worldwide. Here, we perform a global-scale meta-analysis to investigate the extent of the latitudinal survival gradient. We modeled survival as a function of latitude for the separate northern and southern hemispheres, and considered phylogenetic relationships and extrinsic (climate) and intrinsic (life history) predictors hypothesized to moderate these effects. Using a database of 1,004 estimates from 246 studies of avian survival, we demonstrate that in general a latitudinal survival gradient exists in the northern hemisphere, is dampened or absent for southern hemisphere species, and that survival rates of passerine birds largely account for these trends. We found no indication that the extrinsic climate factors were better predictors of survival than latitude alone, but including species’ intrinsic traits improved model predictions. Notably, species with smaller clutch size and larger body mass showed higher survival. Our results illustrate that while some tropical birds may be longer lived than their temperate counterparts, the shape of the latitude-survival gradient differs by geographic region and is strongly influenced by species’ intrinsic traits.

## INTRODUCTION

Aves, a class represented by around 10,000 species, display a broad diversity of morphologies and behaviors, and also show considerable variation in their lifespan and annual survival. For example, in large-bodied landbirds, such as some raptors and parrots, annual survival is often high (Newton *et al*. 2016; Maestri *et al*. 2017) and individuals are long lived, but for small-bodied species like warblers and kinglets, rates of annual survival can be low (DeSante *et al*. 2015; Johnston *et al*. 2016). While differences in body mass account for some of this variation — larger species tend to live longer than smaller ones (Lindstedt & Calder 1976, 1981; Promislow 1993; Speakman 2005) ― many species live longer or shorter lives than predicted given their body mass (Healy *et al*. 2014). Other aspects of a species’ life history, particularly the demographic cost of reproduction, may explain this residual variation in survival rates (Williams 1966; Stearns 1992; Roff 2002). This view stems from the hypothesis that limited resources (i.e., time and/or energy) result in an allocation trade-off between two competing vital functions; specifically, current reproduction reduces future reproduction and survival. The pivotal survival-reproduction trade-off has been well documented in birds (Ricklefs 1977, 2000; Saether 1988; Linden & Møller 1989; Martin 1995; Ghalambor & Martin 2001), and with the observations of early investigators that the number of eggs laid declines from the poles towards the equator (Moreau 1944; Lack 1947; Skutch. 1949), it has given rise to the expectation that tropical species should offset a reduced clutch size by having higher adult survival (Murray 1985).

There are many studies that suggest high adult survival in tropical birds, the majority of which focus on comparisons between north-temperate systems and the tropics. Early reports of high survival came from studies equating survival estimates with return rates (Snow 1962; Fogden 1972; Fry 1980; Bell 1982; Dowsett 1985). While these studies deepened our understanding of life-history strategies in tropical birds, survival-rate estimates based on return rates are problematic because they confound estimation of complicated functions of survival rate and capture probability (Nichols & Pollock 1983; Krementz *et al*. 1989; Sandercock 2006). More recently, studies employing improved methods for estimating survival via Jolly-Seber (JS) and Cormack-Jolly-Seber (CJS) models, which separate apparent survival (i.e., Φ: the product of true survival and site fidelity) from encounter probability (Sandercock 2006), have reinforced the idea of higher adult survival at lower latitudes (Faaborg & Arendt 1995; Johnston *et al*. 1997; Francis *et al*. 1999; Peach *et al*. 2001; McGregor *et al*. 2007). The generality of these findings, however, has been questioned based on comparisons showing negligible differences in survival between Central and North American birds (Karr *et al*. 1990), and lower than expected survival rates for birds from South America (Blake & Loiselle 2008). Other studies have even found higher survival rates for south temperate birds compared to tropical species in Africa (Lloyd *et al*. 2014). Only one quantitative review has formally addressed latitudinal patterns in adult survival rates of birds using survival estimates derived across a broad range of latitudes. Munoz et al. (2018) showed that adult survival was higher for species in the tropics compared to those in five sites across the north temperate zone, supporting the hypothesis of a latitudinal gradient in survival, at least for forest-dwelling passerines in the western Hemisphere. Yet, despite longstanding interest in the idea of a latitudinal gradient in survival, we still lack an empirical synthesis at the global scale, which stands as a limiting factor in our ability to generalize these relationships to the diverse life history of birds found worldwide (Martin 2004).

Most explanations for a latitudinal survival gradient are based on the assumption of consistent latitudinal variation in survival and other life history traits with which it covaries, such as clutch size (Karr *et al*. 1990; Faaborg & Arendt 1995; Johnston *et al*. 1997; Peach *et al*. 2001; McGregor *et al*. 2007). Indeed, most comparative studies of variation in life history traits treat northern and southern latitudes equivalently (Jetz *et al*. 2008; Muñoz *et al*. 2018; Terrill 2018). However, this assumption may not always be met, since latitude itself does not directly influence organisms per se; rather, environmental factors that covary with latitude (i.e., temperature, precipitation, seasonality) exert selective pressures on life history traits. For example, although there exists a global latitudinal gradient in clutch size (Cardillo 2002; Jetz *et al*. 2008), this trend is dampened in the southern hemisphere―south temperate species lay smaller clutches than those in the north temperate hemisphere (Yom-Tov *et al*. 1994; Martin 1996; Evans *et al*. 2005). Consistent with this pattern, south temperate birds in Africa also tend to be longer lived than their north temperate European counterparts (Lloyd *et al*. 2014). This hemispheric asymmetry may in part be due to differing climatic conditions between northern latitudes and equivalent southern ones. Namely, south temperate latitudes are less seasonal and have higher minimum winter temperatures, both of which have been hypothesized to decrease adult mortality and lead to smaller clutch size (Ricklefs 1980; Martin 2004). Similarly, clades and their intrinsic traits that may influence survival rates are also distributed nonrandomly across environmental gradients (Jetz *et al*. 2008; Sibly *et al*. 2012). Migratory habit, for instance, arises at least in part from species occupying higher latitudes and experiencing seasonal environments with lower minimum winter temperatures, and there can be substantial deleterious effects on survival over the migratory phase of the annual cycle (Sillett & Holmes 2002; Rockwell *et al*. 2017). Thus, the geographic variation in survival rates reflects a composite of extrinsic factors, intrinsic traits, and historical events related to a species’ lineage.

Because previous analyses of the latitudinal gradient in survival have focused on the north-temperate / tropical model (Martin 2004; Muñoz *et al*. 2018) and have relied on a narrow group of taxa, our current perspective of the biological underpinnings of the geographic variation in survival rates remains somewhat limited. Here, we present data on survival rates for 679 species of landbirds gathered from across the globe (Fig. 1). The purpose of our meta-analysis was to test the relative importance of latitude and extrinsic climate factors (temperature, precipitation, and seasonality) in explaining geographic patterns of avian survival rates, and to ask whether including intrinsic traits (body mass, clutch size, migratory habit) improved model predictions. Specifically, we ask: (1) Is there a latitudinal gradient in adult survival and, if so, are there differences between hemispheres? (2) Do climate measurements (extrinsic factors) explain differences in survival rates as well as latitude? (3) Do intrinsic traits explain additional variation in species-level survival rates? We tested for these relationships in both nonpasserines and passerines and between Old World and New World birds from island and mainland populations. By integrating data on macroecological processes with comparative biology, our modeling approach provides a powerful tool for understanding the diversity of life histories that have evolved across the globe.

**Figure 1.**
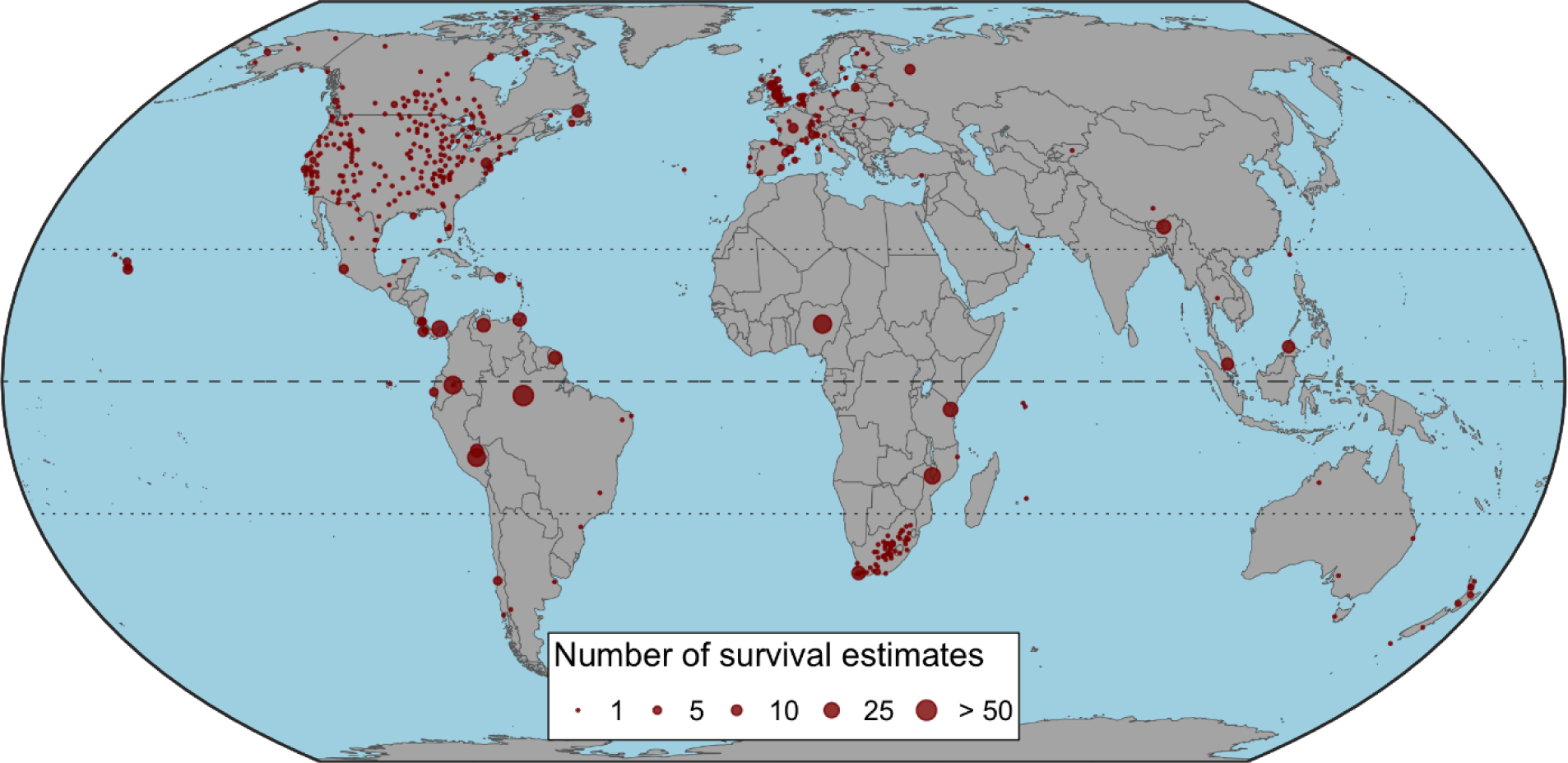
Location of effect sizes from 246 studies used in the meta-analysis of avian survival rates. The number of survival estimates reported at each location is illustrated by the size of the circle. Dashed line represents the equator while dotted lines at 23.4**°** N and S indicate the Tropic of Cancer and Capricorn, respectively, and delineate the tropics.

## METHODS

### Assembling a global dataset of avian survival rates

We conducted an extensive search of the peer-reviewed literature for studies that measured survival rate in bird populations, relying primarily on Web of Science Core Collections (http://apps.webofknowledge.com/) and Google Scholar (https://scholar.google.com/). We also included data for survival rates of North American birds downloaded from the Monitoring Avian Productivity and Survivorship (MAPS) program (www.vitalratesofnorthamericanlandbirds.org; DeSante *et al*. 2015). We searched online using combinations of the following terms: ‘survival’, ‘mortality’, ‘vital rate’, or ‘demography’, and ‘bird’ or ‘avian’. Our initial survey resulted in over 2000 publications. We then screened titles and abstracts of publications and considered them for final inclusion in the meta-analysis if they met the following criteria: (1) the study provided estimates of adult survival, not juvenile or nestling, at the species level. (2) The species studied was not pelagic. (3) The study did not include harvested or captive-bred populations, whose survival rates may not reflect those experienced under natural conditions. (4) Survival rate was estimated on the breeding grounds (i.e., we did not include estimates from studies of over-wintering or migratory stopover sites). (5) The data were collected from marked populations of birds to estimate apparent or true survival using one of three methods: live-recovery/resight models, dead-recovery models, or more complex models that used a combination of these two approaches. (6) The study was conducted for at least 3 years, which is the minimum number of occasions needed to estimate the probability of encounter (*p*) from live-encounter data (Sandercock 2006). To avoid sex-biased differences in survival probability (Székely et al. 2014), we also required that (7) the estimate of survival included data from both males and females. In cases when studies had overlapping data, we retained the publication that presented the most information (i.e., had more precise estimates or used a larger dataset). This second review narrowed the window of appropriate publications to 319. However, many of these papers did not report measurement error on survival rate, which is required to weight individual effect sizes in meta-analyses. Our final dataset therefore consisted of 246 publications (Appendix 1), which we examined in detail and extracted relevant data.

For each publication, we extracted information on species’ annual survival rates and their standard error. Some studies (e.g., Collingham *et al*. 2014) presented standard error of logit survival. In these cases, we used a first order Taylor expansion to approximate precision. When the same study provided separate estimates for males and females, or where estimates were made for different time periods, ages of adult birds, or circumstances (e.g., successful breeders vs. unsuccessful, brood parasites present vs. absent) we took the geometric mean of those estimates. When estimates were available from different habitat types within the same study (e.g., logged vs. unlogged forest), we took the geometric mean of those estimates, provided that the study found no significant differences between groups. If group estimates were reported as significantly different, we chose the estimate from the control group for our analysis. We also recorded information on publication identity, species identity and the method used to estimate adult survival.

### Calculating the latitudinal gradient and global climate data

In order to assess the relationship between survival and latitude, we recorded the geographic coordinates for each species in each study from information provided in the paper itself or by locating the study area on Google Maps (google.com/maps). For 26 studies that measured survival over broad spatial scales, such as at the national or continental level (e.g., the MAPS dataset; DeSante *et al*. 2015), we calculated the centroid of the breeding range for each species within the area specified by the study with occurrence data extracted from eBird using the *R* package *auk* (Strimas-Mackey *et al*. 2018). This allowed us to estimate a unique latitude and longitude for the centroid of each species’ realized breeding range rather than simply selecting an unweighted point in the study area itself. For each target region, we extracted eBird observations submitted over the last 15 years during the breeding period (May 1–August 8 for birds from Canada, the United States, and Europe, and September 1–December 8 for species from South Africa). We further filtered data to include only complete eBird checklists (i.e. those where users specify that all species seen or heard are reported), which allowed us to identify implicit non-detections for each species, and by stationary counts <24 hours, traveling counts <25 km, and area counts within a circle of radius 25 km. We then overlaid the eBird checklists with a 25 km equal-area raster grid and calculated the frequency of occurrence of each species on checklists within each grid cell. We took a weighted average of the grid-cell centers, using the species observation frequency as the weight, and used the geographic coordinates of this centroid to represent the species-site specific latitude. Because latitude is often used as a surrogate for variation in climatic conditions between the north and south poles, we also evaluated the predictive power of three key extrinsic factors that characterize the environment of a species and are hypothesized to influence survival rates: annual precipitation (Wolfe et al. 2015, Shorgen et al. 2019), minimum winter temperature (Robinson *et al*. 2007; Salewski *et al*. 2013), and seasonality (Ricklefs 1980, Martin 2004). We downloaded information on climate measurements from WorldClim Global Climate Data (worldclim.org) averaged over 1970-2000 (Fick and Hijmans 2017) at 2.5-minute spatial resolution. We extracted data for total annual precipitation (*Precip*, mm), minimum temperature of the coldest month (*Temp*_Winter_, ℃) and calculated temperature seasonality (*Temp*_Seasonality_, ℃) following Jetz et al. (2008) as the difference between mean summer and winter temperatures averaged over 3-month periods.

### Clutch size, body mass, and migratory behavior at a global scale

To test additional hypotheses of intrinsic factors that may explain global patterns in avian survival rates, we collected data on body mass, clutch size, and species’ migratory habit. We obtained body mass (measured in grams) from information contained in the paper, or from the “Elton traits” database (Wilman *et al*. 2014), or Handbook of birds of the World Alive (del Hoyo *et al*. 2018). Similarly, we extracted data on clutch size when it was presented in the paper itself and used published standard reference databases (Jetz *et al*. 2008; del Hoyo *et al*. 2018) when it was not. Following del Hoyo *et al*. (2018), we classified species as either migratory or non-migratory. We considered migratory species to be those that regularly undertook seasonal movements >100 km (i.e., short- and long-distance migrants). While some species do not clearly conform to this dichotomy, it is useful way to contrast important sources of mortality that could influence survival rates.

### Statistical analysis

Before conducting analyses, we log10 transformed mass and clutch size due to skewness, and scaled latitude and climate data to z scores by subtracting their mean and dividing by their standard deviation. Most variables were weakly correlated, although both measures of temperature reached Spearman rank correlations >0.75 (Table S1). To estimate adult survival rates along the latitudinal gradient, we used a multi-level meta-analytical framework using the *metafor* package (Viechtbauer 2010), which fits random and fixed effects models, weighting effect sizes by the inverse of their squared standard error. For each model developed, we accounted for sources of non-independence in our dataset that can arise when multiple survival estimates are extracted from the same study, are available for the same species, and / or due to common ancestry, by fitting publication identity, species identity, and phylogeny as random intercepts. To incorporate phylogeny, we used a majority rules consensus tree derived from a set of 1,000 randomly-selected trees based on the global phylogeny of birds (Jetz *et al*. 2012), and pruned to our study taxa (Fig. S1) with the package *phytools* (Revell 2012). We then used the branch length from this consensus tree to specify values for the model variance-covariance matrix. We performed all statistical analyses in *R* environment 3.5.0 (R Core Team 2019).

**Table 1.** Integrated models of survival rates across an assemblage of 679 bird species based on only extrinsic (seasonality and latitude) or intrinsic (body mass, clutch size, and migratory habit) moderators or a combined extrinsic / intrinsic model (Joint model). *Latitude* was fitted with separate intercepts for the Northern and Southern hemisphere. ΔAIC_C_ columns represent the increase in model AIC_C_ when a moderator is dropped relative to the fully parameterized model. Model coefficients (β), 95% confidence intervals are shown for the full models. The Null model AIC for random effects only (phylogeny, study ID, and species nested within study) is listed for comparison.

| Variable | Extrinsic | Intrinsic | Separate models |  |  |  |  | Joint model |  |  |  |  |  |
| --- | --- | --- | --- | --- | --- | --- | --- | --- | --- | --- | --- | --- | --- |
| | $\Delta AIC_C$ | $\Delta AIC_C$ | $\beta$ | 95%<br>LCL | 95%<br>UCL | Z | p | $\Delta AIC_C$ | B | 95%<br>LCL | 95%<br>UCL | z | p |
| *Latitude*<sub>Northern</sub> | 5.84 |  | -0.06 | -0.10 | -0.03 | -3.46 | *** | -2.16 | -0.04 | -0.07 | -0.01 | -2.28 | * |
| *Latitude*<sub>Southern</sub> |  |  | 0.00 | -0.03 | 0.02 | -0.23 |  |  | 0.00 | -0.03 | 0.02 | -0.29 |  |
| *Temp*<sub>Seasonality</sub> | -2.60 |  | -0.01 | -0.03 | 0.01 | -1.09 |  | -4.07 | 0.00 | -0.02 | 0.01 | -0.12 |  |
| *Body mass* |  | 82.97 | 0.05 | 0.04 | 0.06 | 10.04 | *** | 89.12 | 0.05 | 0.05 | 0.07 | 10.35 | *** |
| *Clutch size* |  | 70.99 | -0.12 | -0.14 | -0.09 | -9.29 | *** | 44.97 | -0.10 | -0.13 | -0.07 | -7.40 | *** |
| *Nonmigrant* |  | 4.73 | 0.03 | 0.01 | 0.05 | 3.02 | ** | -2.44 | 0.02 | -0.01 | 0.04 | 1.48 |  |
| AIC<sub>C</sub> of random effects only model = -1518.94 |  |  |  |  |  |  |  |  |  |  |  |  |  |

### Meta-analyses and meta-regressions

We first ran a random effects only model on the entire dataset using the *rma.mv* function to estimate a pooled effect size of global avian survival rates. Given potential differences in selection pressures experienced by nonpasserines vs. passerines, species from Old World (Afrotropics, Indomalayan, Palearctic) vs. New World (Neotropics, Nearctic) biogeographic realms (Olson *et al*. 2001), and island vs. mainland bird populations, we also evaluated separate meta-analytic models using effect sizes for these six data subsets. We considered point estimates to be different from one another if their 95% confidence intervals (CI) did not overlap. We quantified total heterogeneity of each dataset by calculating Cochran’s *Q* and the *I^2^* statistic (Higgins & Thompson 2002).

We conducted meta-regressions (meta-analyses incorporating explanatory variables, hereafter referred to as “moderators”) whereby we determined the effects on species-specific adult survival rates of (1) latitude, (2) extrinsic climatic factors, and (3) intrinsic traits in accordance with hypotheses described from the primary literature. We began by comparing fit of a latitude-only model (*Latitude*), where regression slopes varied between hemispheres, to single-predictor linear models testing the influence of moderators on adult survival rates (Table S2). We next used AIC_C_ values (AIC corrected for small sample size, Burnham & Anderson 2002) to guide selection of a multi-predictor model of extrinsic climatic factors and intrinsic traits separately, and then combined both sets of moderators with *Latitude* into a joint model. Starting with the moderator that had the lowest AIC_C_ value, we sequentially added the next strongest moderator until AIC_C_ was no longer improved. We considered the model that minimized AIC_C_ the most appropriate if it had fewer parameters and was at least 2 AIC_C_ less than the next most competitive model (Arnold 2010). All of the intrinsic moderators we assessed improved model fit (Table S3) and were carried forward to the next step of model development. *Temp*seasonality provided the best model fit for extrinsic moderators; neither annual precipitation nor winter temperature improved AIC_C_ values during construction of the multi-predictor extrinsic model. We repeated analysis of the joint extrinsic / intrinsic model for each of the six data subsets.

### Sensitivity analyses and publication bias

To test whether our results were sensitive to differences in the study method used to estimate survival, we fit additional models with study type as a moderator. We also repeated analyses comparing separate datasets for effect sizes where we used package *auk* to calculate the geographic coordinates versus those where we did not, and again using only those studies that reported survival estimates for less than 10 species, and those that were conducted for a minimum of 10 years. We tested for publication bias of the entire dataset, which can result from selective publication of favorable or statistically significant results, by visually assessing asymmetry of funnel plots.

## RESULTS

Our final dataset consisted of 1004 effect sizes that we extracted from 246 publications for 679 species (Fig. 1; Appendix S1). The majority of effect sizes (82%) came from passerine birds, and effect sizes obtained from the New World biogeographical realms (282 Nearctic and 297 Neotropical) were more numerous than those from Old World studies (176, 169, and 51 of effect sizes from the Palearctic, Afrotropical, and Indomalayan realms, respectively; Fig. S2). Approximately twice as many estimates were available from studies conducted in the northern hemisphere (n = 681) compared to the southern hemisphere (n = 323). The majority of effect sizes reported per study was 1 (geometric mean = 1.6, *SD* = 12.4) and multiple estimates for the same species were available from different studies for 168 species, accounting for 49% of all effect sizes included in the analysis.

### Meta-analytic means and the relationships between intrinsic and extrinsic variables

The meta-analytic mean calculated for the full dataset was 0.66 (95% CI = 0.48 to 0.85; Table S4). This represents the overall global mean survival rate for all birds included in our analysis. The joint model explained variation in survival well (Fig. S3; *x*_Observed_ = 0.40 + 0.38 (residual standard error = 0.07) *x*_Predicted_; *F*_1,1004_ = 738.6; adjusted *r*^2^ = 0.42). When we estimated separate meta-analytical means for the six data subsets, we found similar values with overlapping 95% confidence intervals between the global mean and mean effect sizes calculated for nonpasserines vs. passerines, Old World vs. New World biogeographical realms, and estimates from islands vs. mainland birds (Fig. 2; Table S4). In addition, all models had values of *P*<0.0001 for Q_E_ and *I^2^*>90%, which indicated that substantial heterogeneity remained unexplained among studies and warranted our subsequent step of evaluating moderator variables.

**Figure 2.**
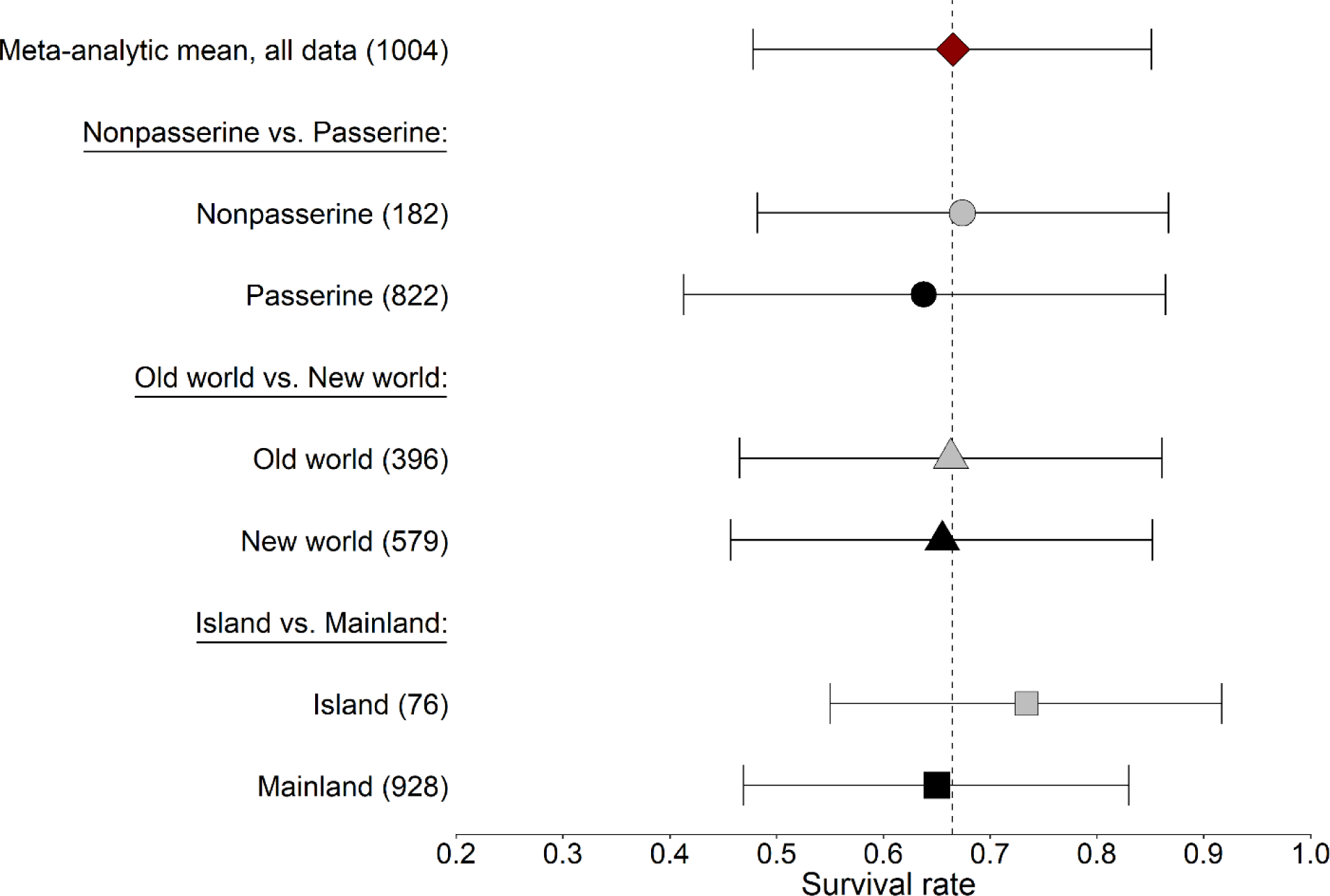
Mean avian survival and 95% confidence limits calculated over the entire dataset and from meta-regression models, which estimated intercepts independently for data from nonpasserine vs. passerine birds, Old World vs. New World biogeographic realms, and islands vs. mainland. Number of effect sizes used in each data subset are shown in parentheses. Dashed line indicates the difference from the overall meta-analytical mean.

We found evidence supporting the hypothesis of a latitudinal gradient in survival, and this effect was most apparent in the northern hemisphere (Figs 3 and 4). When we examined model predictions from a single-predictor model of latitude over the entire dataset, survival decreased by 2% for every 10° increase in latitude for species in the northern hemisphere compared to a <1% decrease for southern hemisphere species (Fig. 3A). Similarly, estimates from the joint model based on the entire dataset showed a significant negative effect of latitude on survival for northern hemisphere species (*β* = −0.039, 95% CI = −0.072 to −0.006), but not for those inhabiting the southern hemisphere (*β* = −0.004, 95% CI = −0.026 to 0.019; Table 1). Driving this global trend at northern latitudes were significant negative point-estimates for passerine birds (*β* = −0.064, 95% CI = −0.103 to −0.025) and species / populations from the mainland (*β* = −0.046, 95% CI = −0.080 to −0.012; Fig. 4). In contrast, effect sizes calculated for southern latitudes were smaller and the overall slope of the meta-regression line of the global model was shallower compared to the northern hemisphere (Figs 3A and 4). Only New World species (i.e., birds from South America) showed a significant negative association with latitude (*β* = −0.055, 95% CI = −0.090 to −0.021; Fig. 4). Of the extrinsic climate moderators we considered, temperature seasonality was the most competitive within our AIC model selection framework (Table S2 and S3), although only marginally so compared to minimum winter temperature. Regardless of which climate moderator was used in the joint model, the effect calculated over the global dataset was nonsignificant (Fig. 3B) and all other data subsets had effect sizes close to and confidence intervals overlapping zero.

**Figure 3.**
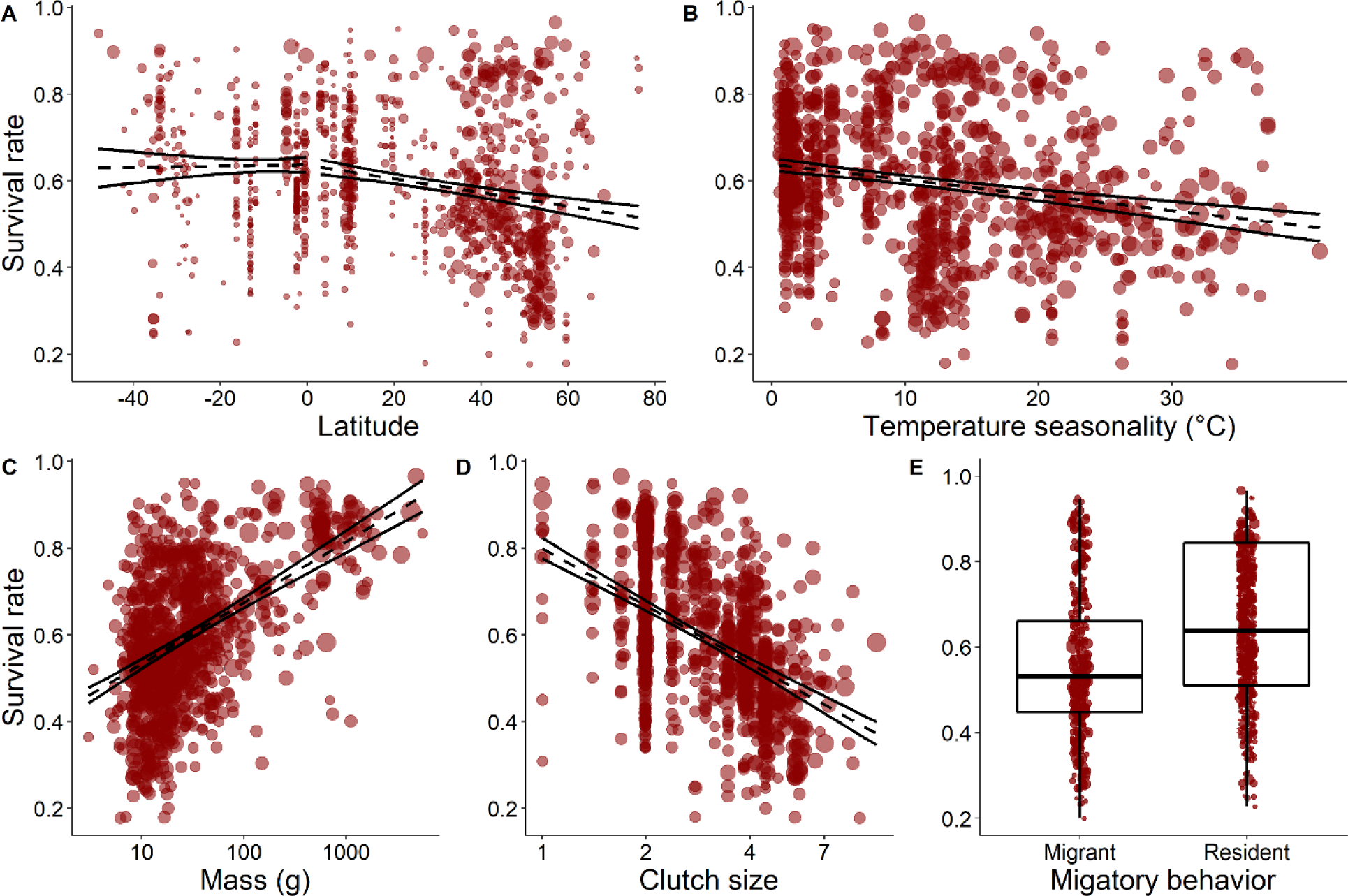
Relationship between adult survival rate of birds from the entire dataset and moderator variables included in the joint extrinsic / intrinsic model (Table 1). Dashed lines represent the best linear fit based on model predictions estimated from single-predictor meta-regression models in *metafor* with 95% confidence intervals plotted as solid lines. Point sizes reflect the inverse of the standard error used to weight data points (i.e., more precise estimates appear as larger points).

**Figure 4.**
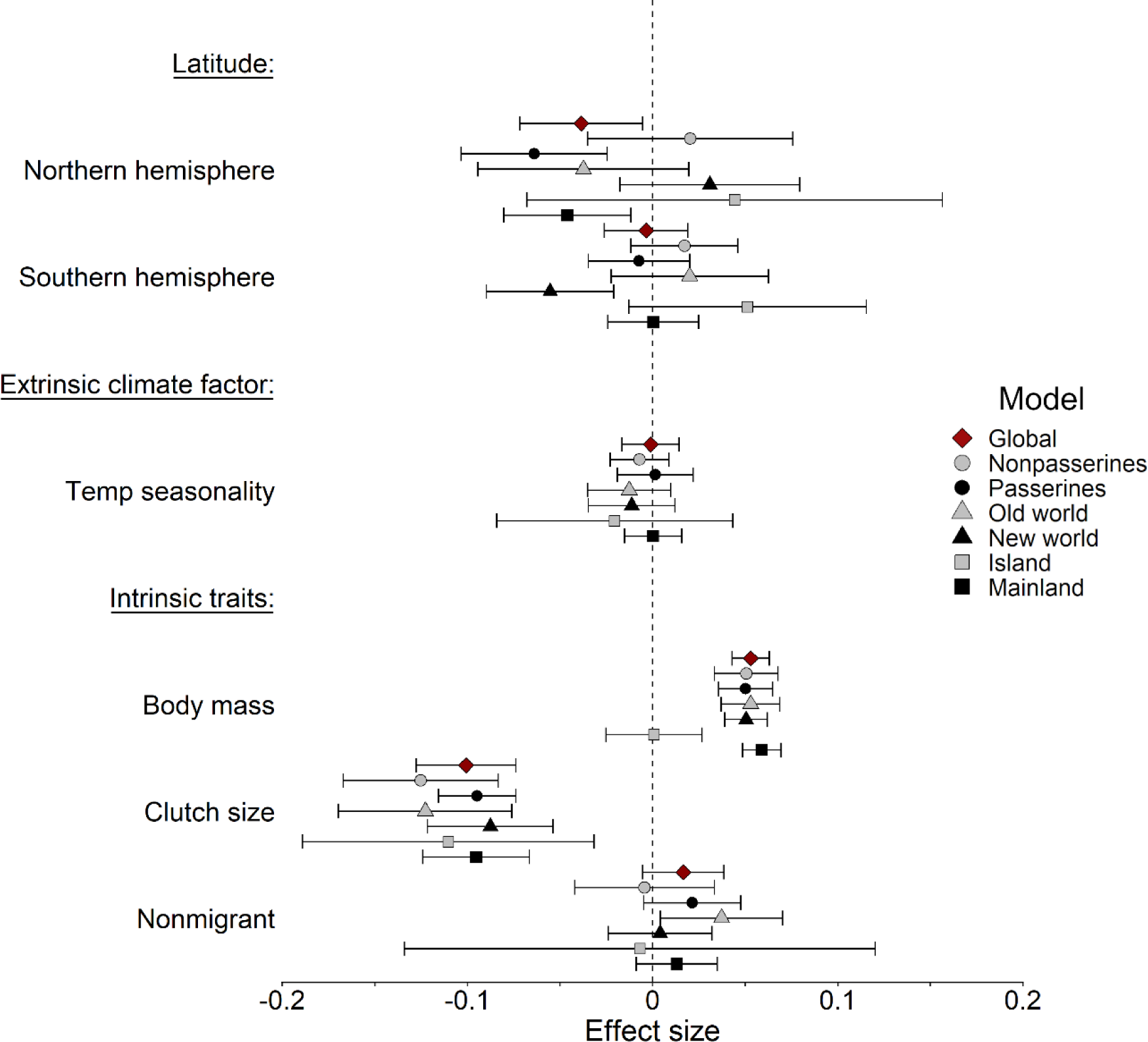
Overall effect sizes for the six variables considered in the joint extrinsic / intrinsic model for the entire dataset and all data subsets. Bars indicate 95% confidence limits. Effect sizes are considered significant where confidence limits do not overlap zero (dashed line).

In general, the relationship between survival and intrinsic life history characteristics was stronger than those of either climate or latitude (Table 1; Fig. 4). Effect size estimated from the global model was positive for mass (*β* = 0.053, 95% CI = 0.043 to 0.063) and negative for clutch size (*β* = −0.101, 95% CI = −0.128 to −0.074), which means that avian survival was higher for larger birds and for those with smaller clutch sizes (Figs 4C and D). With the exception of mass for island species, similar results for both moderators were found for all data subsets, supporting the previously well documented relationship between mass and clutch size and survival (Fig. 4). When we included migration as a moderator in the meta-regression on the full data set, the effect size was small and positive, with confidence intervals marginally overlapping zero (*β* = 0.016, 95% CI = −0.005 to 0.039; Fig. 3E). Although the relative effect of migration varied some between data subsets, all but Old World birds (*β* = 0.037, 95% CI = 0.004 to 0.070) had confidence intervals that included zero, suggesting higher survival for nonmigratory birds in Africa and Asia (Fig. 4).

### Sensitivity analysis

When we estimated mean survival for effect sizes calculated from studies using capture-mark-recapture methods, dead recovery methods, or complex models that integrated a combination of these two approaches, we found point estimates were not distinguishable from the overall mean (Fig. 5). Similarly, when we removed data where we used the package *auk* to calculate latitude and longitude, or when we removed the 22 studies reporting estimates for >10 species (which accounted for nearly 64% of all effect sizes) the results were qualitatively similar to the global mean survival rate based on the entire dataset (Fig. 5). We found no indication of publication bias after examining symmetry of funnel plots (Fig. S4).

**Figure 5.**
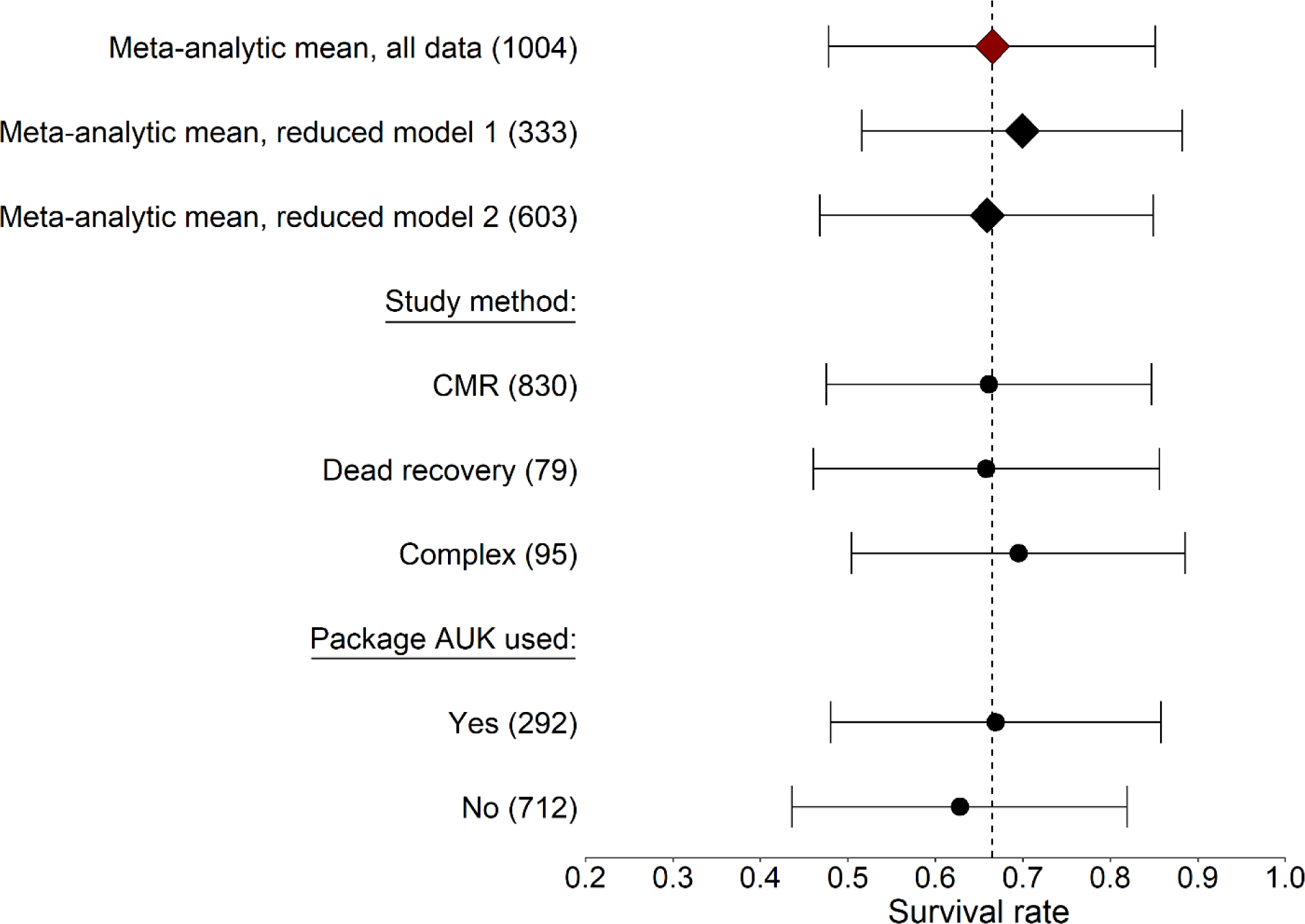
Results of the sensitivity analysis comparing the meta-analytic mean of the entire dataset to a reduced dataset consisting of studies that reported ≤10 effect sizes (reduced model 1), a reduced dataset consisting of studies that collected data for ≥10 years (reduced model 2), effect size estimated separately for each of three methods used to estimate survival, and effect size calculated from studies where we used the package *auk* to approximate species’ geographic coordinates compared to those studies where authors provided information on latitude and longitude. Bars represent 95% confidence limits. Dashed line indicates the difference from the overall meta-analytical mean.

## DISCUSSION

### Global-scale patterns of avian survival with latitude

We found support for the oft-toted latitudinal-survival gradient, but this depended on both the geographic area and taxa being considered. Specifically, we demonstrate that the previously noted inverse relationship between latitude and survival is borne out across northern hemisphere avifauna overall, and that this effect is strengthened when considering only passerines or species inhabiting the mainland (Fig. 4). In contrast, the relationship was only evident in the southern hemisphere for survival estimates from New World birds (Fig. 4), the vast majority of which were passerines. When considered independently, there was no indication that nonpasserines had higher survival with decreasing latitude in either hemisphere. Overall, our meta-analysis reveals that while some tropical birds may be longer lived than their temperate counterparts, the shape of the latitude-survival response is likely to differ among species and between hemispheres.

Our synthesis is the first to assess global-scale patterns in avian survival rates; previous studies have either been limited geographically (Karr *et al*. 1990; Peach *et al*. 2001; Lloyd *et al*. 2014), or have focused on a narrower range of species, such as raptors (Newton *et al*. 2016) or shorebirds (Méndez *et al*. 2018). To date, the most extensive analysis of avian survival and latitude comes from a study of 12 locations spanning 60° across the Americas (Muñoz *et al*. 2018). Our global-scale analysis compliments that of Muñoz *et al*. (2018), who reported a linear decrease in survival of roughly 2.1% for every 10° increase in latitude for passerine birds from Alaska to Peru, nearly identical to what we observed for northern hemisphere species worldwide. Granted both our studies used a meta-analytical approach, Muñoz *et al*. (2018) conducted their analysis using a Bayesian mode of inference and considered only forest-dwelling passerines, while our study includes survival estimates of both passerines and nonpasserines from a variety of habitats, which we investigated using a maximum-likelihood approach. We also fit regression lines for latitude both north and south of the equator rather than testing the relationship between survival and absolute latitude. This latter point is particularly important, given that one general explanation for spatial patterns in life-history traits is that they arise from natural selection imposed by latitudinal gradients in environmental conditions (Cardillo 2002), which differ between hemispheres (Chow et al. 2004). Despite our use of different methods, the fact that we obtained some common results lends increased support to the overall relationship. Moreover, with our analysis, we provide a stronger mechanistic basis for understanding variation in survival rates, as it better reflects the climatic variables that underlie latitude in the northern and southern hemispheres.

Hemispheric asymmetries in other patterns of avian life-history traits, such as timing of reproduction (Covas *et al*. 1999), clutch size (Moreau 1944; Martin *et al*. 2006; Lloyd *et al*. 2014), and parental care (Russell 2000; Russell *et al*. 2004; Llambías *et al*. 2015), are well documented. The global patterns we identified are also congruent with the idea of a differential response of life-histories between hemispheres ― we detected an inverse relationship between survival and latitude in the northern hemisphere but found little indication that this association was mirrored by southern hemisphere species overall. Only when we analyzed biogeographic realms in the southern hemisphere separately did we find that New World birds showed higher survival with decreasing latitude. This pattern is deceptive, however, since southern hemisphere nonpasserines account for little more than 1% of the effect sizes analyzed in the New World data subset. We therefore interpret this result as evidence of the latitudinal-survival gradient in South American passerines. This means that for Old World birds, tropical species had similar survival rates to birds from the austral zone, and this was likely to be true regardless of whether they were passerines or nonpasserines. Survival estimates from Australasia and Oceania, biogeographic realms not traditionally included in the New / Old world classification, also reflected this same pattern and showed no evidence of a negative relationship with latitude.

Such differences may be explained, in part, by the historical geography and latitudinal positions of the continents. For the last 15 million years, South America has extended roughly 20° further into the southern hemisphere than continental landmasses in the Old World. Thus, one reason we may have detected a negative trend in survival for southern hemisphere birds, but only in the New World, could simply be due to the greater range of latitudes and climatic conditions available to landbirds from South America with which to adapt. For example, latitudes greater than 35° S are characterized by higher seasonality and mean annual temperatures ≤0°C (Chown et al. 2004); thus, this result may be indicative of a threshold response of avian survival to freezing temperatures and / or a more seasonal environment. Supporting this idea, mean survival of South American passerines that occurred at latitudes higher than 35° S (survival rate = 0.38, n = 8) was lower on average than those from the highest latitudes occupied by birds in Africa (Old World survival at 34° S = 0.69, n = 19). Only one other study has addressed the question of a latitudinal-survival gradient in the southern hemisphere; Lloyd et al. (2014) found no indication of higher survival for birds living in tropical Malawi compared to austral South Africa. Our results are congruent with those findings and suggest that higher survival of tropical birds may be a pattern localized primarily to passerines from the northern hemisphere and in South America, where factors such as a more seasonal environment may limit resource availability and constrain species survival.

### Influence of Climate on Survival

Our results suggest that temperature seasonality, at least at the resolution at which we examined it, is a poor predictor of avian survival. Indeed, latitude-only models out performed single-predictor models of extrinsic climate factors for each of the moderators we considered by a minimum of >8 ΔAIC_C_ (Table S2). Although temperature seasonality was not significant, our finding of higher survival in the southern hemisphere, but only for New World birds is in accordance with reported asymmetries in climate between hemispheres. Compared to north-temperate latitudes, austral latitudes are characterized as less seasonal in general, having higher minimum winter temperatures and higher, less variable patterns of precipitation (Chown *et al*. 2004). That said, South America does posses environments with climates closer to those of the northern hemisphere (e.g., mean temperatures ≤0°C, higher temperature seasonality) compared to Africa and Asia, which lack such climate analogs at their southern-most latitudes. Although latitudinal variation in life history traits arises in part from natural selection imposed by complex interactions among environmental factors, latitude as a ‘catch-all’ variable provided a more complete picture of global variation in survival. For example, temperature seasonality fails to capture the negative latitude-survival relationship in passerines because this effect is counter-acted by pooling data for taxa from different regions; specifically, combining data with estimates for southern hemisphere passerines from the Old World. It appears, therefore, that latitude remains one of the best methods to portray the suite of climatic constraints that characterize a species’ environment and leads to variation in life histories, but only when northern and southern hemispheres are examined independently.

### Intrinsic traits mediate variation in the latitudinal survival gradient

We find that the association between body mass and survival and reproduction and survival ― two of the cornerstone trade-offs of life history theory (Stearns 1992) ― are well supported by our meta-analysis, suggesting higher survival for larger birds and those with smaller clutch sizes (Fig 3C and D). Notably, when mass and clutch size were included in the joint model, the strength of the latitudinal survival gradient was diminished (Table 1; Fig 3). In contrast, we found the addition of migratory habit to be generally negligible (Fig 3). These results highlight the importance of considering the interplay between intrinsic and extrinsic variables when investigating macroecological processes. Latitude of course influences many aspects of avian life history, including both clutch size (Cardillo 2002; Jetz *et al*. 2008) and body mass (Olson *et al*. 2009), both of which have been demonstrated to increase globally with increasing latitude. Additionally, median body size also decreases systematically within species-rich communities, such as those characterized at many tropical latitudes (Olson *et al*. 2009). Combined with these findings, our results are in accordance with the theory of a slow-fast life-history continuum (Ricklefs & Wikelski 2002) and suggest that while birds at tropical latitudes tend to be longer lived and have reduced clutches given their body size, this is far from the full picture. Global patterns of avian survival are driven by interactions between intrinsic traits and lineage-specific effects of latitude and their associated climatic factors.

### Challenges in evaluating avian survival

Adult survival estimates are affected by several methodological caveats that we consider here. First, a general problem with comparing survival studies is that differences between estimates derived from old versus new methods and between live recaptures and dead recoveries may mask trends in the data (Roodbergen *et al*. 2012). Our dataset consisted primarily of studies that used live capture-mark-recapture techniques (83% of effect sizes) and most of these were conducted since 2000; nearly all studies were conducted after 1990 when modern statistical tools for analyses of marked animals were developed (Lebreton *et al*. 1992). One of the drawbacks of capture-mark-recapture data is that the reported metric, apparent survival, is a product of true survival and site fidelity and as such will always be biased low, whereas estimates of survival from dead recovery models are often interpreted as true survival (Sandercock 2006). Biases in survival estimates may therefore be strong for birds from tropical regions, which consisted exclusively of live-recapture data, and where behaviors such as altitudinal migration are more common than in temperate regions (Barçante *et al*. 2017) and can lead to permanent emigration from study plots. Another issue affecting the comparison of survival studies is the study duration. This, too, may be particularly problematic for tropical regions, where data collection is often hampered by sampling conducted over irregular or insufficiently long intervals to produce robust estimates of survival (Ruiz-Guitérrez *et al*. 2012). For example, in our meta-analysis 62% and 69% of effect sizes from austral and temperate latitudes, respectively, were calculated from datasets spanning >10 years, compared to only 34% from tropical latitudes. However, in a study of tropical birds comparing survival estimates derived from 6 vs.12 years of data, Blake & Loiselle (2013) reported an improvement in precision, but no change in point estimates for survival. Still, other authors argue that longer time frames are needed to generate reliable survival estimates for tropical resident species (i.e., 10–30 years), given their expected longevity and low recapture probabilities (*p* <0.25; Ruiz-Guitérrez *et al*. 2012). Despite these problems with the comparability of the data, we found no indication that difference in methodological approaches strongly biased our results (Fig. 5).

## CONCLUSION

Based on a global-scale synthesis of avian survival rates, we find evidence that survival increases with decreasing latitude, but that this phenomenon is more nuanced than previous descriptions have characterized. Specifically, we demonstrate that the latitudinal survival gradient is stronger in northern hemisphere species, where climate seasonality may be greater. By including aspects of species life history characteristics in our models, we could explain a greater portion of the variation in survival rates than with latitude alone. These results indicate the importance of considering an organism’s intrinsic traits as well as the extrinsic factors of their environment when describing broad-scale macroecological patterns. Where peaks in survival occur, how they relate to climatic variables, and how these patterns are likely to change through time and space given the effects of climate change, are of major importance for conservation. We hope that in assembling this database and dissecting some of the global patterns in survival across avian groups and hemispheres, we can provide a platform for future work to target underrepresented regions and taxa and also make a clear path forward to better understanding variation in survival rates, and how it intersects with other life history traits across the world’s avifauna.

## ACKNOWLEDGEMENTS

We thank Peter Arcese and Ben Freeman for comments on previous versions of this manuscript. We would also like to thank Umesh Srinivasan for helping locate studies of avian survival throughout the world. This research was supported by funding from the University of British Columbia to M. N. Scholer, and by Discovery Grant F11-05365 from the Natural Sciences and Engineering Research Council to J. E. Jankowski.

---

**Figure S1.**
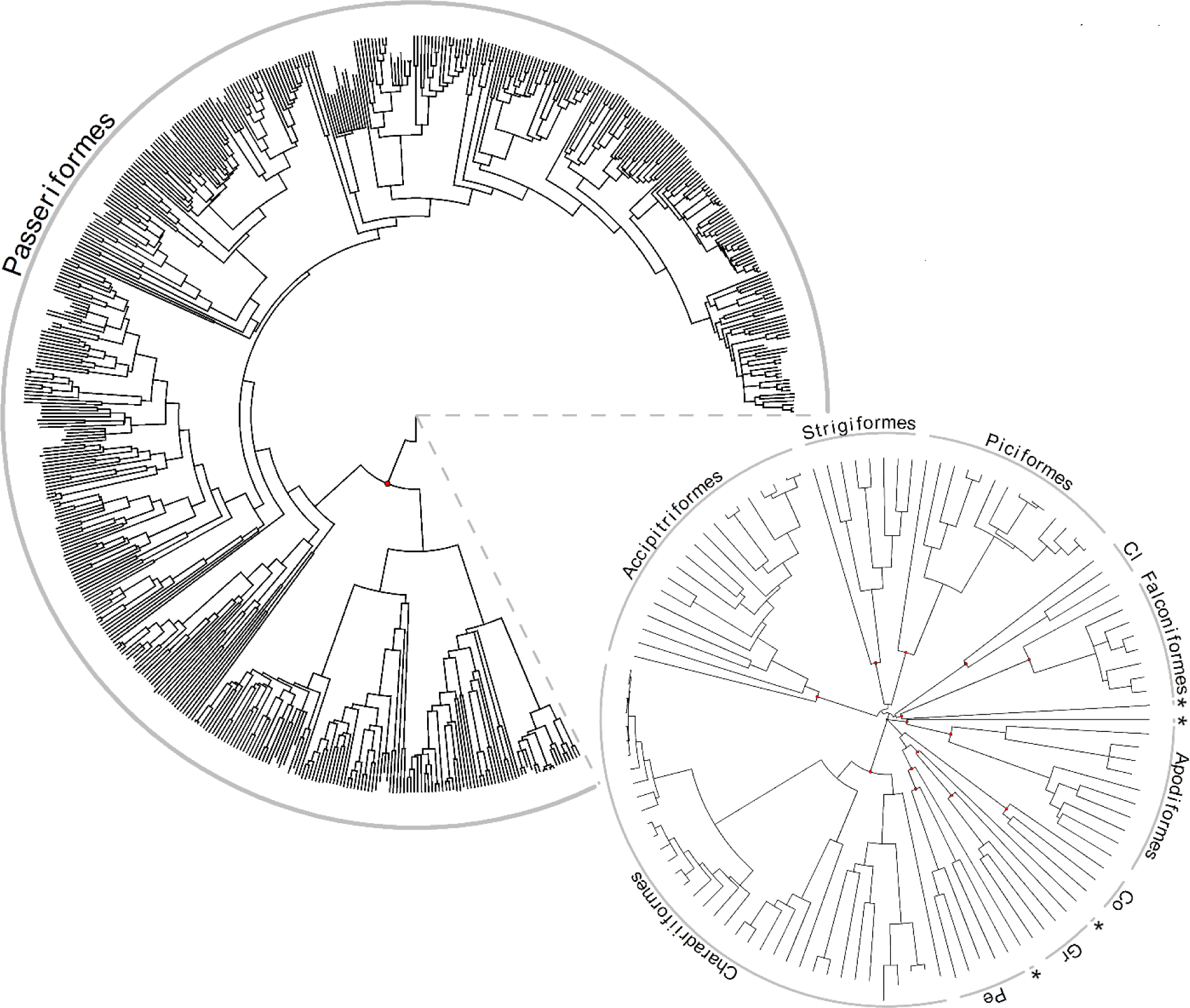
Phylogenetic relationships of the 679 species of birds used in the global meta-analysis of survival rates. Most species (82%) were in order Passeriformes. The call out shows non-passerine orders used in the analysis. Red circles indicate nodes demarking branches for each of the orders. Abbreviations are: Cl = Coliformes, Co = Columbiformes, Gr = Gruiformes, Pe = Pelcaniformes. From clockwise from top right, asterix symbols show orders represented by a single species: Psittasciformes, Caprimulgiformes, Cuculiformes, Ciconiformes.

**Figure S2.**
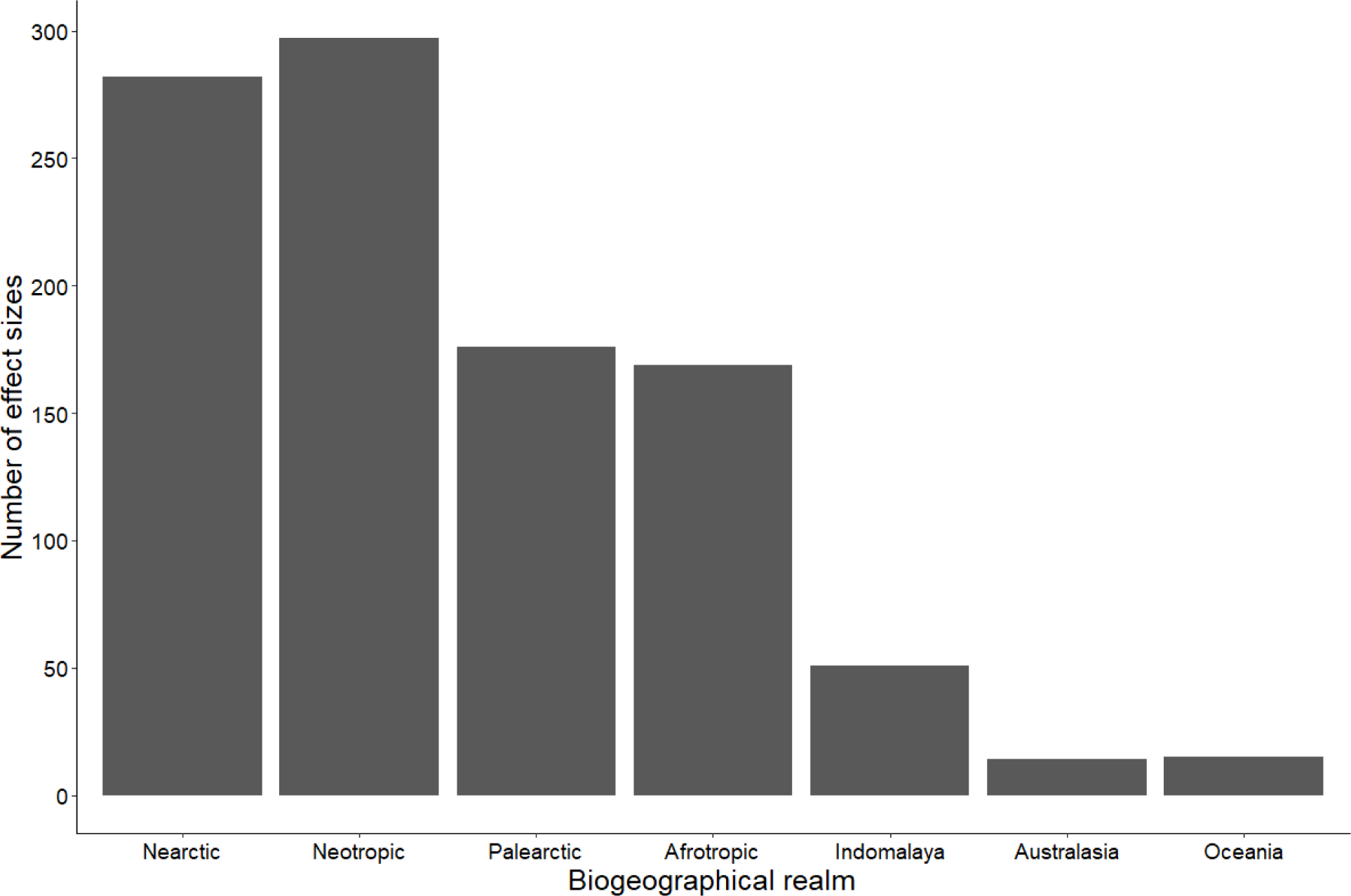
Number of effect sizes coming from the biogeographic realms of the world excluding the Antarctic.

**Figure S3.**
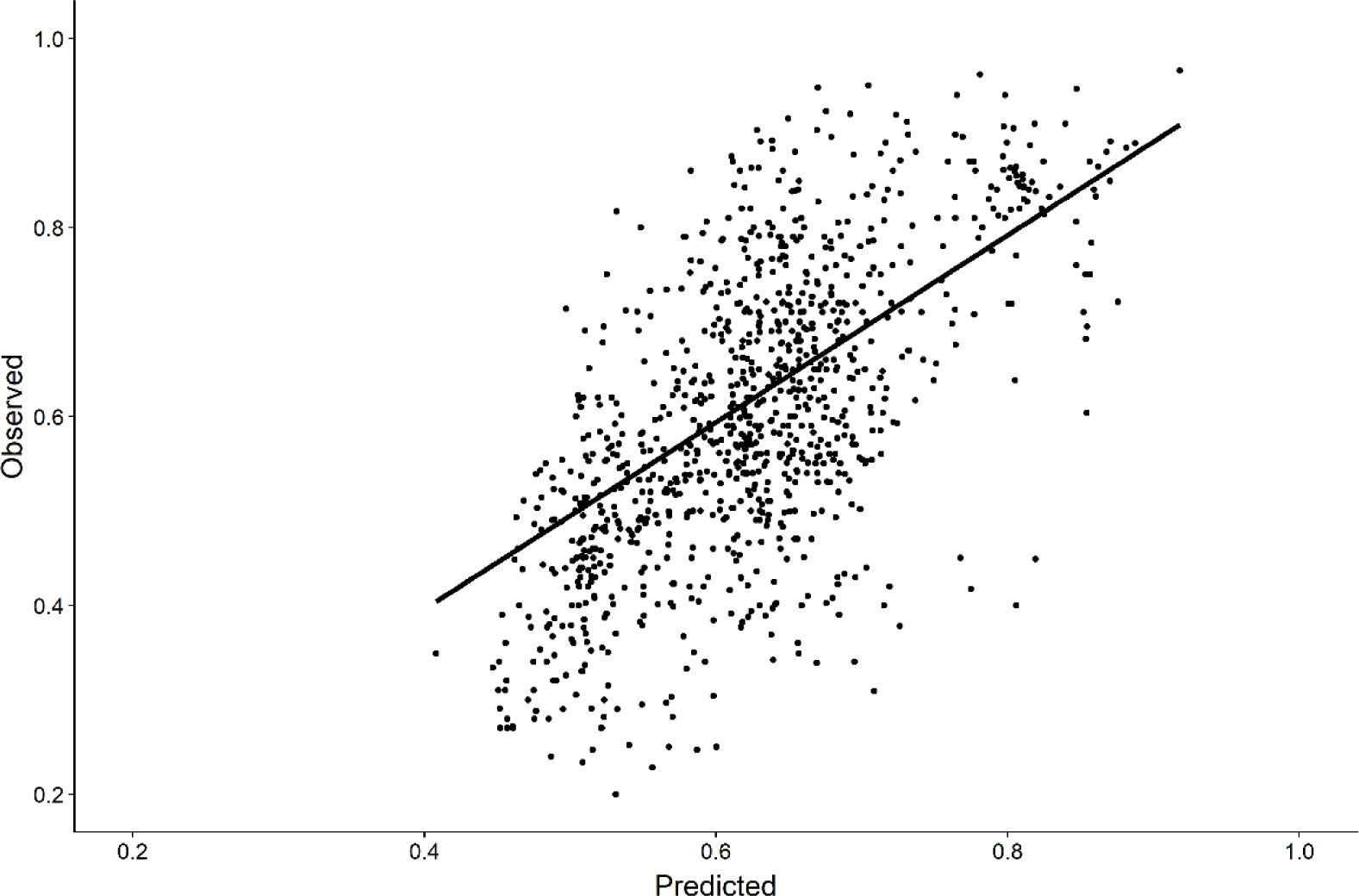
Model fit for the observed survival rate and that predicted by the joint extrinsic / intrinsic model (Table 1) for an assemblage of 679 species of birds across the globe.

**Figure S4.**
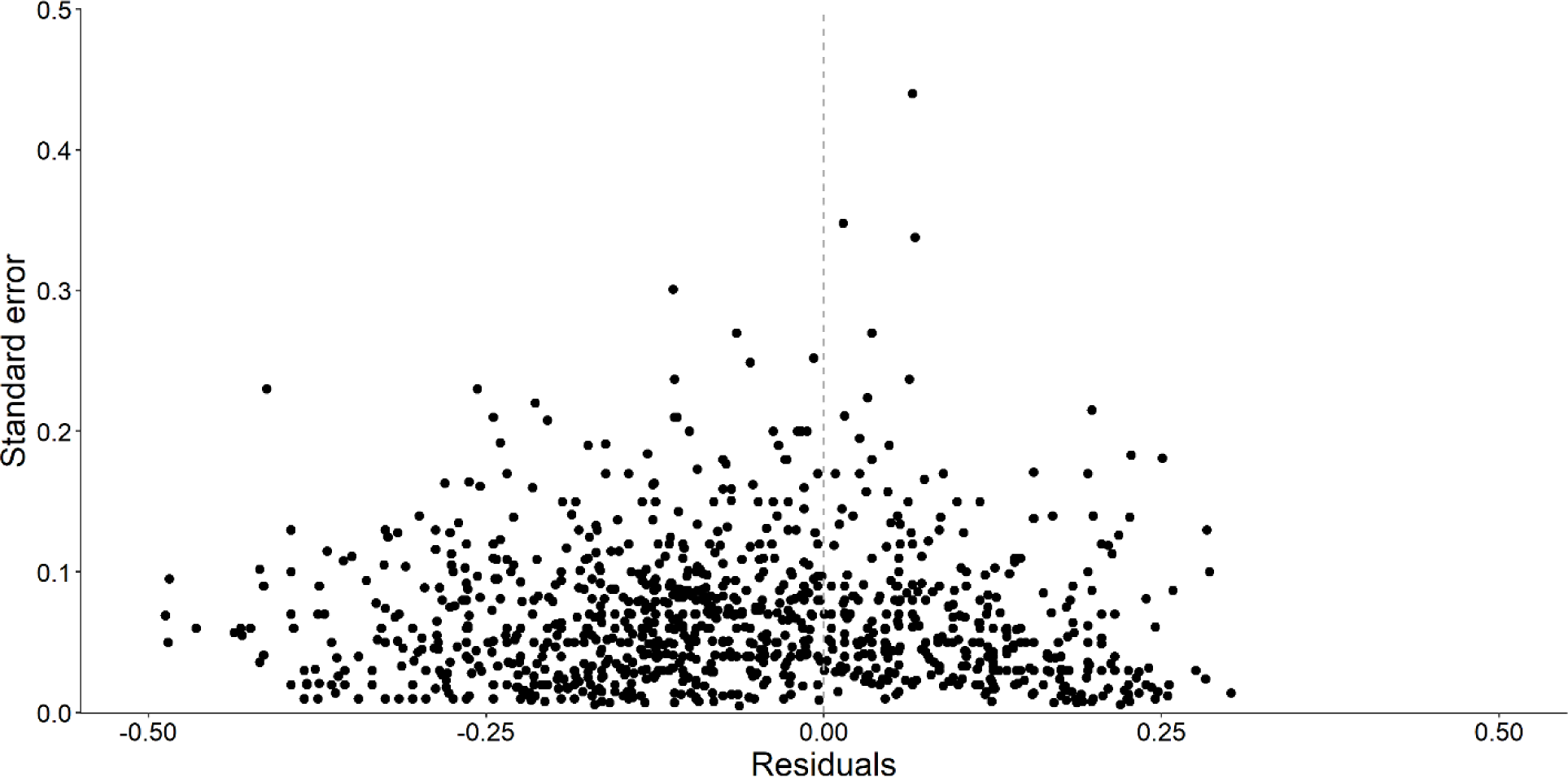
Funnel plot used to evaluate publication bias for the global analysis of 1004 effect sizes plotted against their precision.

**Table S1.**
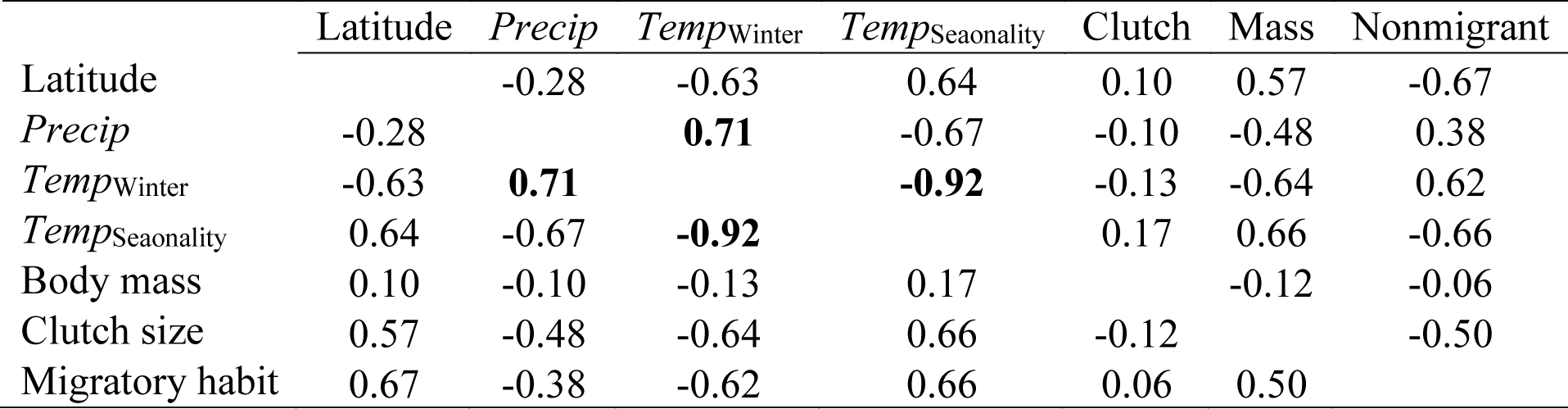
Spearman rank correlations of variables considered for the meta-analysis of avian survival rates. Significant correlations ≥ 0.70 are highlighted in bold.

|  | Latitude | *Precip* | *Temp*<sub>Winter</sub> | *Temp*<sub>Seasonality</sub> | Clutch | Mass | Nonmigrant |
| --- | --- | --- | --- | --- | --- | --- | --- |
| Latitude |  | -0.28 | -0.63 | 0.64 | 0.10 | 0.57 | -0.67 |
| *Precip* | -0.28 |  | **0.71** | -0.67 | -0.10 | -0.48 | 0.38 |
| *Temp*<sub>Winter</sub> | -0.63 | **0.71** |  | **-0.92** | -0.13 | -0.64 | 0.62 |
| *Temp*<sub>Seasonality</sub> | 0.64 | -0.67 | **-0.92** |  | 0.17 | 0.66 | -0.66 |
| Body mass | 0.10 | -0.10 | -0.13 | 0.17 |  | -0.12 | -0.06 |
| Clutch size | 0.57 | -0.48 | -0.64 | 0.66 | -0.12 |  | -0.50 |
| Migratory habit | 0.67 | -0.38 | -0.62 | 0.66 | 0.06 | 0.50 |  |

**Table S2.** Single-predictor meta-regression models of extrinsic and intrinsic moderators hypothesized to effect adult survival rates and ranked by AIC_C_.

| Model | *K* | AIC<sub>C</sub> | $\Delta$ AIC<sub>C</sub> |
| --- | --- | --- | --- |
| Mass | 5 | -1606.11 | 0.00 |
| Clutch | 5 | -1600.92 | 5.19 |
| Latitude | 6 | -1540.16 | 65.96 |
| Seasonality | 5 | -1531.7 | 74.41 |
| Winter | 5 | -1531.18 | 74.94 |
| Migratory | 5 | -1527.77 | 78.34 |
| Precipitation | 5 | -1519.29 | 86.82 |
| Null | 4 | -1516.05 | 90.07 |

**Table S3.**
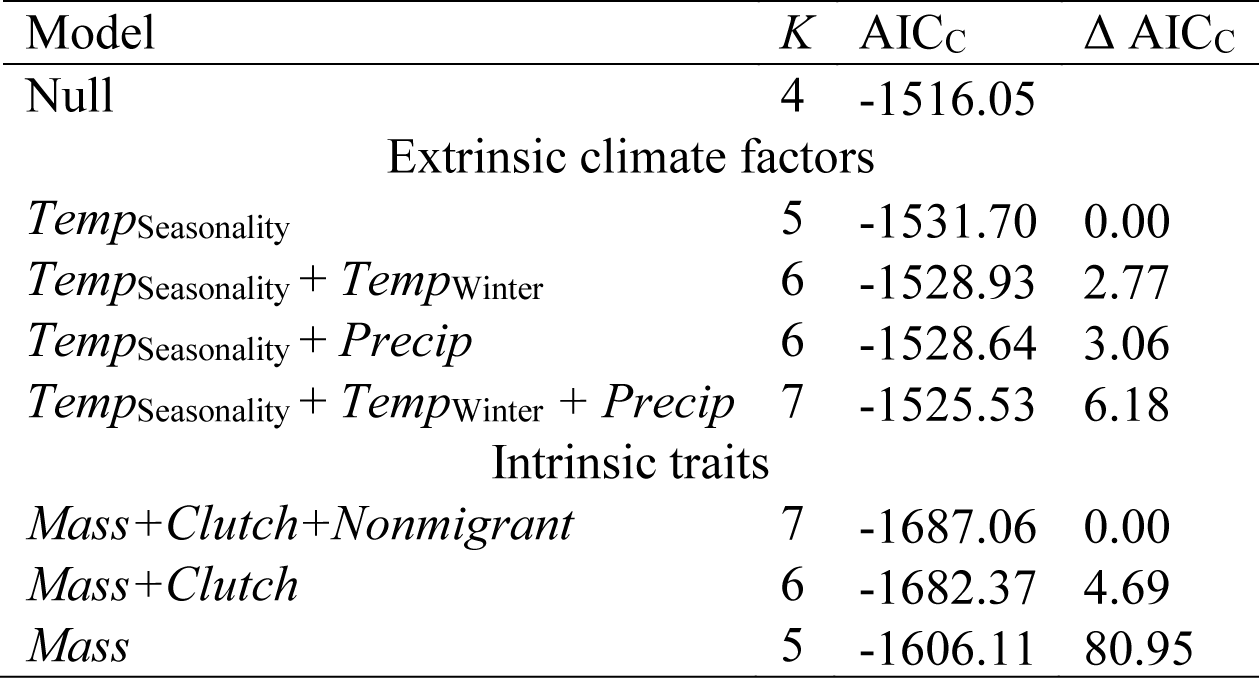
Multi-predictor meta-regression models of either extrinsic climate factors or intrinsic traits. Models were developed by sequentially adding the best performing moderators until AIC was no longer improved, then combining these to form the joint extrinsic / intrinsic model.

**Table S4.** Mean survival rate of each data subset, their upper and lower bound %95 confidence interval, the number effect sizes in the analysis, and two measure of total heterogeneity: Chochran’s *Q* and *I_2_* values.

| Data used | Mean survival | 95% CI lower | 95% CI upper | No. effect sizes used | Cochran's Q | *I*<sup>2</sup> |
| --- | --- | --- | --- | --- | --- | --- |
| All data | 0.66 | 0.48 | .85 | 1004 | *P* > 0.001 | 0.99 |
| Nonpasserines | 0.67 | 0.48 | 0.87 | 182 | *P* > 0.001 | 0.99 |
| Passerines | 0.64 | 0.41 | 0.86 | 822 | *P* > 0.001 | 0.96 |
| Old world | 0.65 | 0.46 | 0.85 | 396 | *P* > 0.001 | 0.96 |
| New world | 0.66 | 0.47 | 0.86 | 579 | *P* > 0.001 | 0.94 |
| Island | 0.73 | 0.55 | 0.92 | 76 | *P* > 0.001 | 0.91 |
| Mainland | 0.65 | 0.47 | 0.83 | 928 | *P* > 0.001 | 0.96 |

## REFERENCES

1. Arnold, T.W. (2010). Uninformative parameters and model selection using Akaike’s information criterion. J. Wildl. Manage., 74, 1175–1178.

2. Barçante, L., M. Vale, M. & Maria, M.A. (2017). Altitudinal migration by birds: A review of the literature and a comprehensive list of species. J. F. Ornithol., 88, 321–335.

3. Bell, H.L. (1982). Survival among birds of the understorey in lowland rainforest in Papua New Guinea. Corella, 6, 77–82.

4. Blake, J.G. & Loiselle, B.A. (2008). Estimates of apparent survival rates for forest birds in eastern Ecuador. Biotropica, 40, 485–493.

5. Blake, J.G. & Loiselle, B.A. (2013). Apparent survival rates of forest birds in eastern Ecuador revisited: Improvement in precision but no change in estimates. PLoS One, 8, 8–13.

6. Burnham, K.P. & Anderson, D.R. (2002). Model selection and multimodel inference: A practical information-theoretic approach. 2nd ed. Springer-Verlag, New York, NY.

7. Cardillo, M. (2002). The life-history basis of latitudinal diversity gradients: How do species traits vary from the poles to the equator? J. Anim. Ecol., 71, 79–87.

8. Chown, S.L., Sinclair, B.J., Leinaas, H.P. & Gaston, K.J. (2004). Hemispheric asymmetries in biodiversity: A serious matter for ecology. PLOS Biol., 2, 1701–1707.

9. Collingham, Y.C., Huntley, B., Altwegg, R., Barnard, P., Beveridge, O.S., Gregory, R.D., et al. (2014). Prediction of mean adult survival rates of southern African birds from demographic and ecological covariates. Ibis (Lond. 1859)., 156, 741–754.

10. Covas, R., Lepage, D., Boix-Hinzen, C. & du Plessis, M. (1999). Evolution of sociality and life-history strategies in birds: Confronting northern perspectives in the southern hemisphere. S. Afr. J. Sci., 95, 400–402.

11. DeSante, D.F., Kaschube, D.R. & Saracco, J.F. (2015). Vital rates of North American Landbirds. Institue Bird Popul. Available at: www.VitalRatesOfNorthAmericanLandbirds.org. Last accessed 1 May 2019.

12. Dowsett, A.R.J. (1985). Site-fidelity and survival rates of some montane forest birds in Malawi south-central Africa. Biotropica, 17, 145–154.

13. Evans, K.L., Duncan, R.P., Blackburd, T.M. & Crick, H.Q.P. (2005). Investigating geographic variation in clutch size using a natural experiment. Funct. Ecol., 19, 616–624.

14. Faaborg, J. & Arendt, W.J. (1995). Survival rates of Puerto Rican birds: Are islands really that different? Auk, 112, 503–507.

15. Fogden, M.P.L. (1972). The seasonality and population dynamics of equatorial forest birds in Sarawak. Ibis (Lond. 1859)., 114, 307–343.

16. Francis, C.M., Terborgh, J.S. & Fitzpatrick, J.W. (1999). Survival rates of understorey forest birds in Peru. In: Proceedings 22nd International Ornithological Congress, Durban, South Africa, 16-22 August 1998 (eds. Adams, N.J. & Slotow, R.H.). BirdLife South Africa, Johannesburg, South Africa, pp. 326-335.

17. Fry, C.H. (1980). Survival and longevity among tropical land birds. In: Proceedings of the 4th Pan-African Ornithilogical Congress, Mahé, Seychelles, 6-13 May 1976 (ed. Johnson, D.N.). Southern African Ornithological Society, Johannesburg, South Africa, pp. 333–343.

18. Ghalambor, C.K. & Martin, T.E. (2001). Fecundity-survival trade-offs and parental risk-taking in birds. Science (80-.)., 292, 494–497.

19. Healy, K., Guillerme, T., Finlay, S., Kane, A., Kelly, S.B.A., McClean, D., et al. (2014). Ecology and mode-of-life explain lifespan variation in birds and mammals. Proceedings. Biol. Sci., 281, 20140298.

20. Higgins, J.P.T. & Thompson, S.G. (2002). Quantifying heterogeneity in a meta-analysis. Stat. Med., 21, 1539–1558.

21. del Hoyo, J., Elliott, A., Sargatal, J., Christie, D. & E, de J. (2018). Handbook of the Birds of the World Alive.

22. Jetz, W., Sekercioglu, C.H. & Böhning-Gaese, K. (2008). The worldwide variation in avian clutch size across species and space. PLOS Biol., 6, e303.

23. Jetz, W., Thomas, G.H., Joy, J.B., Hartmann, K. & Mooers, A.O. (2012). The global diversity of birds in space and time. Nature, 491, 444–448.

24. Johnston, A., Robinson, R.A., Gargallo, G., Julliard, R., van der Jeugd, H. & Baillie, S.R. (2016). Survival of Afro-Palaearctic passerine migrants in western Europe and the impacts of seasonal weather variables. Ibis (Lond. 1859)., 158, 465–480.

25. Johnston, J.P., White, S.A., Peach, W.J. & Gregory, R.D. (1997). Survival rates of tropical and temperate passerines: A Trinidadian perspective. Am. Nat., 150, 771–789.

26. Karr, J.R., Nichols, J.D., Klimkiewicz, M.K. & Brawn, J.D.J.D. (1990). Survival rates of birds of tropical and temperate forests: Will the dogma survive? Am. Nat., 136, 277–291.

27. Krementz, D.G., Sauer, J.R. & Nichols, J.D. (1989). Model-based estimates of annual survival rate are preferable to observed maximum lifespan statistics for use in comparative life-history studies. Oikos, 56, 203–208.

28. Lack, D. (1947). The significance of clutch-size. Ibis (Lond. 1859)., 89, 302–352.

29. Lebreton, J.-D.D., Burnham, K.P., Clobert, J. & Anderson, D.R. (1992). Modeling survival and testing biological hypotheses using marked animals: A unified approach with case studies. Ecol. Monogr., 62, 67–118.

30. Linden, M. & Møller, A.P. (1989). Cost of reproduction and covariation of life history traits in birds. Trends Ecol. Evol., 4, 367–371.

31. Lindstedt, S.L. & Calder, W.A. (1976). Body size and longevity in birds. Condor, 78, 91–94.

32. Lindstedt, S.L. & Calder, W.A. (1981). Body size, physiological time, and longevity of homeothermic animals. Q. Rev. Biol., 56, 1–16.

33. Llambías, P.E., Carro, M.E. & Fernández, G.J. (2015). Latitudinal differences in life-history traits and parental care in northern and southern temperate zone House Wrens. J. Ornithol., 156, 933–942.

34. Lloyd, P., Abadi, F., Altwegg, R. & Martin, T.E. (2014). South temperate birds have higher apparent adult survival than tropical birds in Africa. J. Avian Biol., 45, 493–500.

35. Maestri, M.L., Ferrati, R. & Berkunsky, I. (2017). Evaluating management strategies in the conservation of the critically endangered Blue-throated Macaw (Ara glaucogularis). Ecol. Modell., 361, 74–79.

36. Martin, T.E. (1995). Avian life history evolution in relation to nest sites, nest predation, and food. Ecol. Monogr., 65, 101–127.

37. Martin, T.E. (1996). Life history evolution in tropical and south temperate birds: What do we really know? J. Avian Biol., 27, 263–272.

38. Martin, T.E. (2004). Avian life-history evolution has an eminent past: Does it have a bright future? Auk, 121, 289–301.

39. Martin, T.E., Bassar, R.D., Bassar, S.K., Fontaine, J.J., Lloyd, P., Mathewson, H.A., et al. (2006). Life-history and ecological correlates of geographic variation in egg and clutch mass among passerine species. Evolution (N. Y)., 60, 390–398.

40. McGregor, R., Whittingham, M.J. & Cresswell, W. (2007). Survival rates of tropical birds in Nigeria, West Africa. Ibis (Lond. 1859)., 149, 615–618.

41. McNab, B.K. (1994). Resource use and the survival of land and freshwater vertebrates on oceanic islands. Am. Nat., 144, 643–660.

42. Méndez, V., Alves, J.A., Gill, J.A. & Gunnarsson, T.G. (2018). Patterns and processes in shorebird survival rates: A global review. Ibis (Lond. 1859)., 160, 723–741.

43. Moreau, R.E. (1944). Clutch-size: A comparative study, with special reference to African birds. Ibis (Lond. 1859)., 86, 286–347.

44. Muñoz, A.P., Kéry, M., Martins, P.V. & Ferraz, G. (2018). Age effects on survival of Amazon forest birds and the latitudinal gradient in bird survival. Auk, 135, 299–313.

45. Murray, B.G. (1985). Evolution of clutch size in tropical species of birds. Ornithol. Monogr., 36, 505–519.

46. Newton, I., McGrady, M.J. & Oli, M.K. (2016). A review of survival estimates for raptors and owls. Ibis (Lond. 1859)., 158, 227–248.

47. Nichols, J.D. & Pollock, K.H. (1983). Estimation methodology in contemporary small mammal capture-recapture studies. J. Mammal., 64, 253–260.

48. Olson, D.M., Dinerstein, E., Wikramanayake, E.D., Burgess, N.D., Powell, G.V.N., Underwood, E.C., et al. (2001). Terrestrial ecoregions of the world: A new map of life on earth. Bioscience, 51, 933–938.

49. Olson, V.A., Davies, R.G., Orme, C.D.L., Thomas, G.H., Meiri, S., Blackburn, T.M., et al. (2009). Global biogeography and ecology of body size in birds. Ecol. Lett., 12, 249–259.

50. Peach, W.J., Hanmer, D.B. & Oatley, T.B. (2001). Do southern African songbirds live longer than their European counterparts? Oikos, 93, 235–249.

51. Promislow, D.E.L. (1993). On size and survival: Progress and pitfalls in the allometry of life span. J. Gerontol., 48, B115–B123.

52. R Core Team. (2019). R: A language and environment for statistical computing. Vienna, Austria.

53. Revell, L.J. (2012). An R package for phylogenetic comparative biology (and other things). Methods Ecol. Evol., 3.

54. Ricklefs, R.E. (1977). On the evolution of reproductive strategies in birds: Reproductive effort. Am. Nat., 111, 453–478.

55. Ricklefs, R.E. (1980). Geographical variation in clutch size among passerine birds: Ashmole’s hypothesis. Auk Ornithol. Adv., 97, 38–49.

56. Ricklefs, R.E. (2000). Density dependence, evolutionary optimization, and the diversification of avian life histories. Condor, 102, 9–22.

57. Ricklefs, R.E. & Wikelski, M. (2002). The physiology/life-history nexus. Trends Ecol. Evol., 17, 462–468.

58. Robinson, R.A., Baillie, S.R. & Crick, H.Q.P. (2007). Weather-dependent survival: Implications of climate change for passerine population processes. Ibis (Lond. 1859)., 149, 357–364.

59. Rockwell, S.M., Wunderle, J.M., Sillett, T.S., Bocetti, C.I., Ewert, D.N., Currie, D., et al. (2017). Seasonal survival estimation for a long-distance migratory bird and the influence of winter precipitation. Oecologia, 183, 715–726.

60. Roff, D. (2002). Life history evolution. Sinauer Associates, Sunderland, Massachusetts.

61. Roodbergen, M., van der Werf, B. & Hötker, H. (2012). Revealing the contributions of reproduction and survival to the Europe-wide decline in meadow birds: Review and meta-analysis. J. Ornithol., 153, 53–74.

62. Ruiz-Guitérrez, V., Doherty, P.F., Santana, E.C., Contreras Martínez, S., Schondube, J., Verdugo Munguía, H., et al. (2012). Survival of resident Neotropical birds: Considerations for sampling and analysis based on 20 years of bird-banding efforts in Mexico. Auk, 129, 500– 509.

63. Russell, E.M. (2000). Avian life histories: Is extended parental care the southern secret? Emu, 100, 377–399.

64. Russell, E.M., Yom-Tov, Y. & Geffen, E. (2004). Extended parental care and delayed dispersal: Northern, tropical, and southern passerines compared. Behav. Ecol., 15, 831–838.

65. Saether, B.-E. (1988). Pattern of covariation between life-history traits of European birds. Nature, 331, 616–617.

66. Salewski, V., Hochachka, W.M. & Fiedler, W. (2013). Multiple weather factors affect apparent survival of European passerine birds. PLoS One, 8, e59110–e59110.

67. Sandercock, B.K. (2006). Estimation of demographic parameters from live-encounter data: A summary review. Source J. Wildl. Manag., 70, 1504–1520.

68. Scholer, M.N., Arcese, P., Londoño, G.A., Puterman, M. & Jankowski, J.E. (2019). Data from: Survival is negatively related to basal metabolic rate in tropical Andean birds. Dryad Digit. Repos.

69. Sibly, R.M., Witt, C.C., Wright, N.A., Venditti, C., Jetz, W. & Brown, J.H. (2012). Energetics, lifestyle, and reproduction in birds. Proc. Natl. Acad. Sci., 109, 10937–10941.

70. Sillett, T.S. & Holmes, R.T. (2002). Variation in survivorship of a migratory songbird throughout its annual cycle. J. Anim. Ecol., 71, 296–308.

71. Skutch., A.F. (1949). Do tropical birds rear as many young as they can nourish? Ibis (Lond. 1859)., 91, 430–455.

72. Snow, D.W. (1962). A field study of the Black and White Manakin, Manacus manacus, in Trinidad. Zoologica, 47, 65–104.

73. Speakman, J.R. (2005). Body size, energy metabolism and lifespan. J. Exp. Biol., 208, 1717– 1730.

74. Stearns, S.C. (1992). The evolution of life histories. Oxford Univeristy Press, London.

75. Strimas-Mackey, M., Miller, E. & Hochachka, W. (2018). auk: eBird Data Extraction and Processing with AWK.

76. Terrill, R.S. (2018). Feather growth rate increases with latitude in four species of widespread resident Neotropical birds. Auk, 135, 1055–1063.

77. Viechtbauer, W. (2010). Conducting meta-analyses in R with the metafor package. J. Stat. Softw., 36, 1–48.

78. Williams, G.C. (1966). Natural selection, the costs of reproduction, and a refinement of Lack’s principle. Am. Nat., 100, 687–690.

79. Wilman, H., Belmaker, J., Simpson, J., de la Rosa, C., Rivadeneira, M.M. & Jetz, W. (2014). EltonTraits 1.0: Species-level foraging attributes of the world’s birds and mammals. Ecology, 95, 2027–2027.

80. Yom-Tov, Y., Iglesias, G.J. & Christie, M.I. (1994). Clutch size in passerines of southern South America. Condor Ornithol. Appl., 96, 170–177.

